# Combinatorial engineering reveals shikimate pathway bottlenecks in para-aminobenzoic acid production in *Pseudomonas putida*

**DOI:** 10.1101/2024.06.17.599342

**Authors:** Marco A Campos-Magaña, Sara Moreno-Paz, Vitor AP Martins dos Santos, Luis Garcia-Morales, Maria Suarez-Diez

## Abstract

Combinatorial approaches in metabolic engineering enable the optimization of multigene pathways, thereby improving product titers. However, the optimization of complex metabolic pathways is hindered by their multiple interactions. Testing all possible combinations of suitable genetic parts is often prevented by the large number of possible variants. A valuable alternative to this is to use statistical design of experiments and linear modeling to collect important information for optimization without testing every possible combination. The shikimate pathway is an example of a complex metabolic pathway involved in the production of aromatic compounds, which are prevalent in industry. In this study, we explore the impact of the modulation of the expression levels of all the genes in the shikimate and para-aminobenzoic acid (pABA) biosynthesis pathways for pABA production (a widely used industrial intermediate) in *Pseudomonas putida*. We used this approach to select 14 representative strains from a total of 512 possible combinations. We obtained a range of product titers from 2 to 186.2 mg/l. This information was used to guide a second round of strain construction to further increase the production to 232.1 mg/l. Using this strategy, we demonstrate that *aroB* expression, encoding 3-dehydroquinate synthase, is a significant limiting factor in the production of pABA.

## 1 Introduction

Industrial biotechnology uses microbial cell factories as an alternative to the petroleum-based chemical industry, a sector with the third largest greenhouse gas emissions in industry^1^. Specifically, the shikimate pathway, responsible for the synthesis of aromatic amino acids, is considered a potential source of relevant compounds including polymer precursors (cis,cis-muconic acid), food additives (tryptophan), pharmaceuticals (salicylic acid), aromatic agents (vanillin) and biofuels (2-phenylethanol) currently produced via chemical conversion from petroleum-derived benzene, toluene, and xylene^2,3^ . This metabolic pathway has been exploited for the microbial production of numerous highly valuable compounds. Notably, the ShikiFactory100 project, aims to produce over 100 different compounds derived from this pathway, underscoring its significant potential^4^

The shikimate pathway starts with the condensation of erythrose 4-phosphate and phosphoenolpyruvate to form 3-deoxy-D-arabino-heptulosonate 7-phosphate, which, through a series of six reactions, is converted into chorismate (Figure 1). Chorismate can then be converted into the three aromatic amino acids, phenylalanine, tyrosine, and tryptophan, as well as serve as a precursor for other aromatic compounds ^5^. However, obtaining high titers of aromatic compounds is challenged by the supply of precursors, the presence of regulatory systems, and cytotoxicity ^3^. The tight regulation of the shikimate pathway results in low metabolic fluxes that hinder product formation. Although a common approach to increase the carbon flux towards production is the over-expression of shikimate pathway genes, there is no agreement on which genes to over-express for each specific product or microbial host. For instance, production of the shikimate-derived product protocatechuic acid in *Pseudomonas putida* benefited from the over-expression of a feedback-resistant variant of *aroG* (*aroG*^D146N^) and the over-expression of *aroQ* ^6^. In *Escherichia coli* the over-expression of *aroG*^D146N^ generally benefits the production of shikimate-derived products, over-expression of *tyrA* has shown beneficial effects for the production of coumarins, and salvianic acid, while over-expression of *aroB, aroE* and *aroK* were specific for the production of violacein, salvianic acid, and tyrosine, respectively ^5,7^. Alternatively, shikimate production was limited in *Bacillus subtilis* by the expression of *aroA* and *aroD*, homologs of *aroG* and *aroE* ^8^.

**Figure 1.**
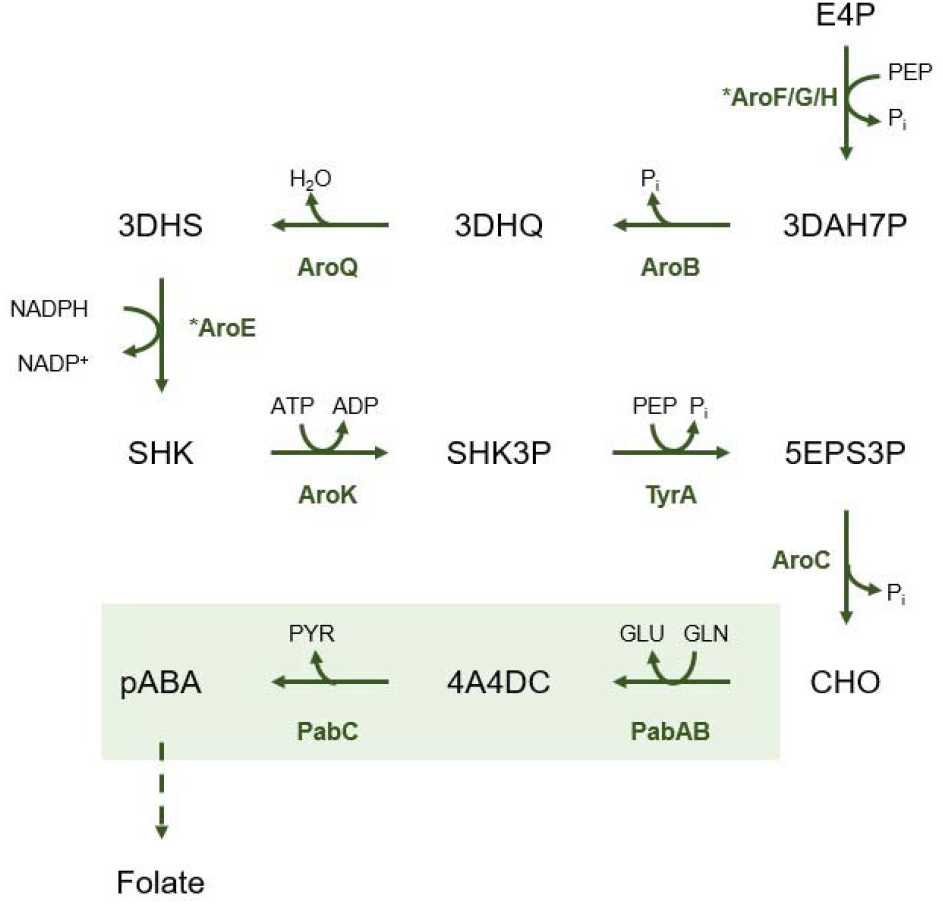
Biosynthetic pathway of pABA. E4P, erythrose 4-phosphate; PEP, phosphoenolpyruvate; P_i_, phosphate; 3DAH7P, 3-deoxyarabino-heptulosonate 7-phophate; 3DHQ, 3-dehydroquinate; 3DHS, 3-dehydroshikimate; SHK, shikimate; SH3P, shikimate 3-phophate; 5EPS3P, 5-enoylpyruvoyl-shikimate 3-phophate; CHO, chorismate; 4A4DC, 4-amino-4-deoxychorismate; pABA, p-aminobenzoic acid. AroG, 3-deoxyarabinoheptulosonate synthase; AroB, 3-dehydroquinate synthase; AroQ, 3-dehydroquinate dehydratase; AroE, quinate/shikimate dehydrogenase; AroK, shikimate kinase; TyrA, 3-phosphoshikimate 1carboxylvinyl transferase; AroC, chorismate synthase; PabAB, aminobenzoate synthase; pabC, 4amino-4-deoxychorismate lyase. Enzymes subject to feed-back regulation are marked with * ^3,5^.

Here we propose the use of statistical design of experiments (DoE) to explore the effect of the expression of shikimate pathway genes on product formation while minimizing the number of strains to construct and test. As opposed to one-factor-at-a-time experimentation, which would require testing the effect of individual gene over-expressions, DoE uses orthogonal designs that allow the identification of the effect of individual genes when they are over-expressed in combinations. In this way, interactions among genes, such as synergistic effects of simultaneous over-expressions can be detected ^9–13^. The genes to over-express in each strain are selected to prevent confounding of individual gene effects. Multiple DoE designs are available^9,13^. Among the different DoE designs, we used Plackett Burman design to explore the impact of the modulation of levels of over-expression of all genes in the shikimate pathway on product accumulation. This design allows identification of partial correlations between individual gene effects and interaction effects^12–14^. Once the strains are constructed, their production data is used to train a linear model in which each factor (i.e. gene) is associated to a model coefficient that determines the effect of the gene on production. Finally, an analysis of variance (ANOVA) is employed to identify genes with a significant positive or negative effect on product titer^9,13^.

We used production of para-aminobenzoic acid (pABA) in *P. putida* as a case study for the application of DoE to identify genes in the shikimate pathway limiting production. Improving the efficiency of the shikimate pathway in *P. putida* is important due to its central role in the biosynthesis of aromatic compounds. *P. putida* has been postulated as a microbial chassis with inherent stress-resistance capabilities and high-levels of NADPH, a cofactor essential for the biosynthesis of shikimate pathway-derived compounds^2,15–17^. In bacteria, pABA is synthesized from chorismate by the action of three enzymes: PabA, PabB, and PabC (Figure 1) ^2^. pABA is a precursor for the formation of folate and it also serves as a precursor for the pharmaceutical industry and as a crosslinking agent for the synthesis of resins and dyes ^18,19^.

## 2 Materials and methods

### 2.1 Promoter and vector backbone selections

Promoter sequences JE111111 and JE151111 and ribosome binding sites JER04 and JER10 were selected from a characterized promoter-ribosome binding site collection ^20^. Plasmids pSEVA231 (medium copy number) and pSEVA621 (low copy number) conferring resistance to kanamycin and gentamycin, respectively, were used for strain construction ^21^. All strains and plasmids used in the present study are listed in Table S1 and S2, respectively. Primers used for plasmid construction are listed in Table S3.

### 2.2 Strain construction

Plasmids were built from individual genetic parts, comprising vector backbone, synthetic promoter, and enzyme coding genes with ribosome binding sites (RBS) and terminator (Figure S1). For low gene expression level, pSEVA621, JE151111, and JER10 were used as backbone, promoter, and RBS, respectively. For the high gene expression level, pSEVA231, JE111111, and JER04 were used as backbone, promoter, and RBS, respectively. All three para-aminobenzoic genes (the complete ORFs) were amplified from genomic DNA of *E. coli* K-12. Shikimate pathway genes including *aroK, aroE, aroB, aroQ, tyrA*, and *aroC* were amplified from genomic DNA of *P. putida* KT2440. *aroG* was amplified from *E. coli* K-12 genomic DNA using primers that introduced a mutation to generate the aroG^D146N^ alelle. All DNA fragments were amplified using Q5® Hot Start High-Fidelity DNA Polymerase (New England Biolabs). DNA fragments were purified with NucleoSpin™ gel and PCR clean-up kits (Macherey-Nagel, Germany). Synthetic promoters were ordered from IDT (Integrated DNA Technologies). DNA assembly of the different constructs was performed using Golden Gate or Gibson assembly. All plasmids were transformed by heat shock in chemically competent *E. coli* DH5_αλ_ pir and selected on LB agar with corresponding antibiotics. Colonies were screened through colony PCR with Phire Hot Start II DNA Polymerase (Thermo Fisher Scientific) using the screening primers listed in the Table S3. Plasmids were verified using Sanger sequencing (Macrogen inc.) or whole plasmid sequencing (Plasmidsaurus inc.) and subsequently transformed into *P. putida* KT2440 via electroporation. For this, 100 ng of plasmid was electroporated into 100 µl cell suspension aliquots with a voltage of 2.5 kV, 25 μF capacitance, and 200 Ω resistance ^22^.

### 2.3 Bacterial strains and growth conditions

*P. putida* KT2440 and *E. coli* cultures were incubated at 30 ºC and 37 ºC respectively. For cloning purposes, both strains were propagated in Lysogeny Broth (LB) medium. For the preparation of solid media, 1.5 % (w/v) agar was added. Antibiotics to select colonies harboring plasmids were used at the following concentrations: kanamycin (Km) 50 µg/ml and gentamycin (Gm) 10 µg/ml. All growth experiments were performed using M9 minimal medium (per liter; 3.88 g K_2_HPO_4_, 1.63 g NaH_2_PO_4_, 2.0 g (NH_4_)_2_SO_4,_ 10 mg ethylenediaminetetraacetic acid (EDTA), 0.1 g MgCl_2_·6H_2_O, 2 mg ZnSO_4_·7H_2_O, 1 mg CaCl_2_·2H_2_O, 5 mg FeSO_4_·7H_2_O, 0.2 mg Na_2_MoO_4_·2H_2_O, 0.2 mg CuSO_4_·5H_2_O, 0.4 mg CoCl_2_·6H_2_O, 1 mg MnCl_2_·2H_2_O, pH 7.0). Strains were precultured overnight in 10 ml LB with corresponding antibiotics. pABA production experiments were performed with 12 ml of M9 minimal medium supplemented with 70 mM glucose and corresponding antibiotics. Cells were grown for 48 h at 30 ºC and 250 rpm in 50 ml mini-bioreactor tubes (Corning) in an Innova 44 incubator (New Brunswick Scientific). At the end of the cultivation, samples for OD_600nm_ measurements and pABA quantification were taken. Production of pABA was measured using three colonies as biological replicates to evaluate variations for each experiment.

### 2.4 Analytical methods

Cell growth was determined by measuring the optical density at 600 nm (OD_600nm_) using an OD600 DiluPhotometer spectrophotometer (IMPLEN). pABA titer was determined using HPLC (Shimadzu) with a C18 column (4.6 mm × 250 mm) and a UV/vis detector set at 235 nm. The mobile phase consisted of Milli-Q water (A), 100 mM formic acid (B), and acetonitrile (C) with a flow rate of 0.75 ml/min at 30 °C. Chromatographic separation of analytes was attained using the following gradient program: t = 0 – 5min: A-80%, B-10%, and C-10%; from t = 5 – 12.5 min: A-0%, B-10% and C-90% and then t = 12.5 – 15 min: A-80%, B-10% and C-10%.

### 2.5 Experimental design and statistical analysis

R (version 4.3.3) was used for the generation of the design and data analysis. Plackett Burman (PB) designs were generated with the ‘pb’ function from the FrF2 R package given the number of factors and experiments^23^. Experimental data was used to train a linear model by least square linear regression using the lm R function. The summary function was used to obtain the ANOVA table which provides the estimated coefficients and their associated p-values that were corrected to account for multiple testing using Bonferroni. The adjusted coefficient of determination (Adj R^2^) was used to assess the model fit to experimental data.

Differences in pABA concentration among experiments were evaluated by two-tailed homoscedastic t-tests.

## 3 Results

### 3.1 Identification of factors affecting pABA production

The synthesis of pABA from erythrose 4-phosphate and phosphoenolpyruvate requires 10 genes involved in nine enzymatic reactions: seven reactions from the shikimate pathway and two committed reactions for pABA production (Figure 1). Therefore, we selected all the genes required for the synthesis of pABA as candidates for over-expression to optimize production. We used *aroB, aroQ, aroE, aroK, aroA*, and *aroC* genes from *P. putida* and the feedback-resistant variant *aroG*^D146N^ from *E. coli* ^24^. Besides, the pABA biosynthetic genes *pabA, pabB*, and *pabC* from *E. coli* K-12 were selected as targets for over-expression.

We used a Plackett Burman design to explore the effect of gene over-expressions on pABA production. In this design combinations of genes are simultaneously over-expressed so the effect of each gene over-expression on pABA production can be determined^12,14^. We considered each of the overexpression target genes as a single factor except for *pabA* and *pabB* which were considered together. The proteins coded by these two genes form a complex and therefore their unbalanced expression was expected to negatively impact pABA production^25^. Each of the factors was explored at two levels including moderate and high over-expression based on copy number plasmid, strength of the promoter, and ribosome binding site assigned to each gene. The high over-expression level was obtained by combining the medium copy number plasmid pSEVAb231 ^21^ with the strong promoter JE111111 and JER04 RBS^20^. A milder over-expression was achieved by combining the low copy number plasmid pSEVAb621 ^21^ with the JE151111 promoter and the JER10 RBS^20^. While studying the combinatorial effect of nine factors and two levels would require the construction of 512 strains (2^9^), the Plackett Burman design reduced the number of strains to build to 16 (2^4^), represented in Figure 2A.

**Figure 2.**
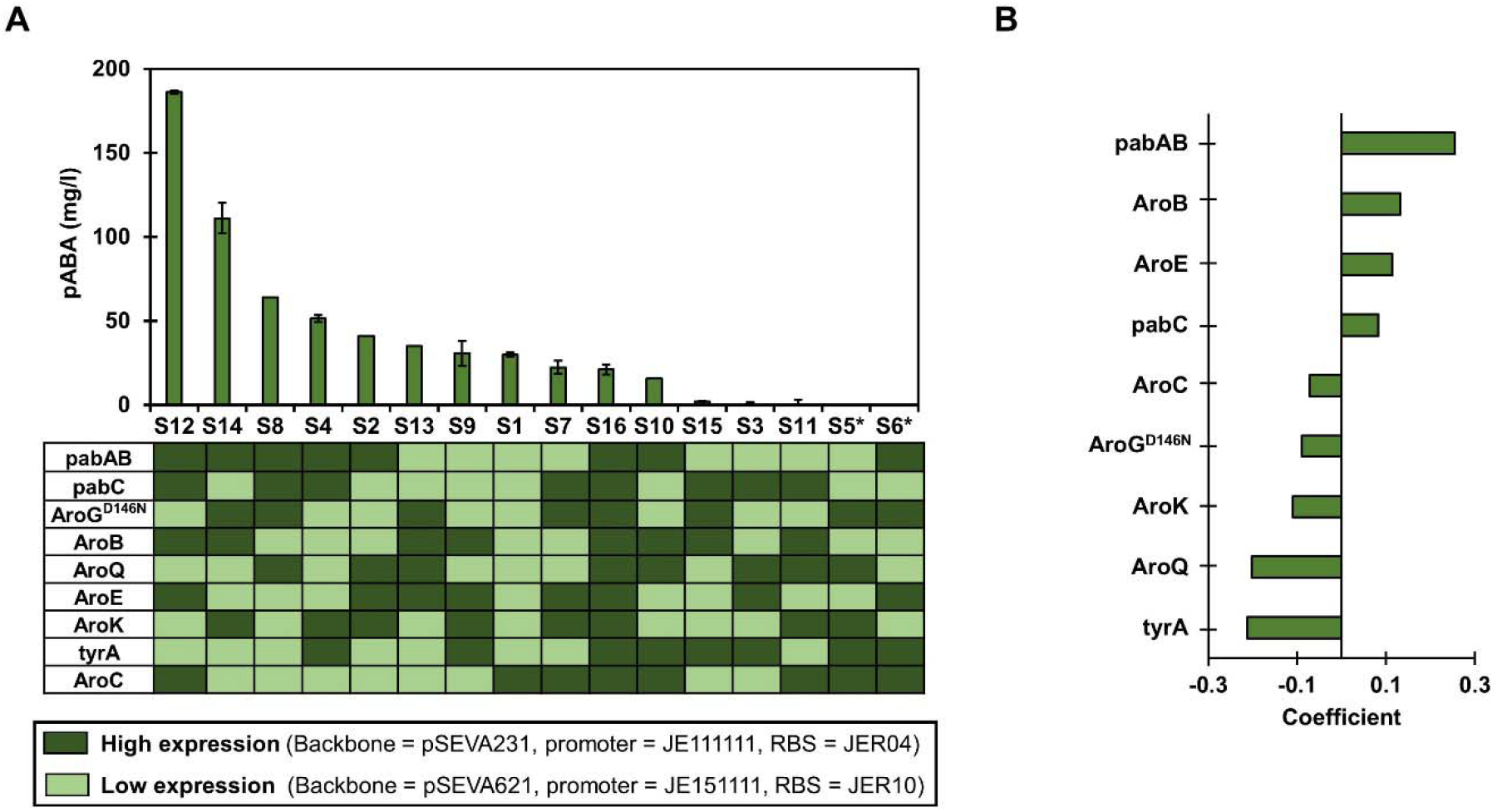
Identification of factors affecting pABA production. **A**. Expression strength of the genes in the strains constructed following a Plackett Burman design (S1 to S16) and their pABA production after 48 h culture in minimal media. Note that strains S5 and S6 could not be constructed. **B**. Regression coefficients of the linear model trained with data from A. Regression coefficients: *pabAB* = 0.25, *aroB* = 0.13, *aroE* = 0.11, *aroC* = 0.08, *tyrA* = -0.21, *aroQ* = -0.20, *aroK= -0*.*10, aroG*^D146N^ = -0.08, and *aroC* = -0.07. All the coefficients are significant according to ANOVA with p-values < 0.05 corrected using Bonferroni.

Fourteen of the sixteen strains required for the design were successfully constructed. Colonies with correct DNA assemblies for strains S5 and S6 were not found (Figure 2A), which did not affect the ability of the design to estimate the effect of all the genes^13^. The pABA production data measured with the available strains was sufficient to train a linear model (Adj R^2^= 0.94) and obtain estimates of the coefficients for each of the tested factors (Figure 2B). Across the different strains, pABA production exhibited a two-order-of-magnitude variation, ranging from 2.0 ± 3.4 mg/l to 186.2 ± 0.32 mg/l, demonstrating the effect of changing the expression levels of the selected genes on pABA production (Figure 2A). The ANOVA on the linear model coefficients reported a significant effect of all the factors on pABA production (Figure 2B). The high over-expression of *pabAB* had the highest positive effect on pABA production, followed by *aroB*, and *aroE* . Therefore, a high expression of the *pabAB, aroB*, and *aroE* genes is essential to obtain high pABA titers. A weaker effect was observed for *pabC* . In addition, we identified genetic factors with negative regression coefficients, and therefore, a negative effect of high over-expression on pABA production. *TyrA* and *aroQ* had the highest negative effect on pABA production. They were followed by *aroK, aroG*^D146N^, and *aroC* (Figure 2B)

Notably, the expression strength levels of the genes in strain S12, the measured strain with the best titer, corresponded with the sign of all the estimated coefficients except for *aroC* (Figure 2). The negative regression coefficients of the linear model suggest that overall high gene over-expression negatively affects pABA production as confirmed by strain S16.

### 3.2 Optimization of pABA production by expanding the design space

After identifying the effect of different degrees of over-expression of the tested genes, we intended to further improve pABA titers by testing a broader range of expression levels. First, the effect of native gene expression was compared to mild over-expression, excluding high over-expression that resulted in low pABA titers. Additionally, we used bicistronic designs as gene expression enhancers for the genes whose over-expression had a positive effect on pABA titers ^26,27^.

High over-expression of *pabAB* was identified as the factor with the highest positive influence on pABA production. Therefore, strains with these genes in the high-expression plasmid were used as background to study the effect of basal expression of the other genes. Additionally, a strain only over-expressing *pabABC* was used as control (C) to evaluate the impact of over-expressing shikimate pathway genes. In all cases, reducing gene expression from low over-expression to basal expression had a significant negative effect on pABA production (Figure 3A), indicating that, even if high over-expression is detrimental, mild over-expression is beneficial for production. The control strain also showed lower pABA titers compared to S12 showcasing that the over-expression of *pabABC* is not enough to obtain high pABA titers.

**Figure 3.**
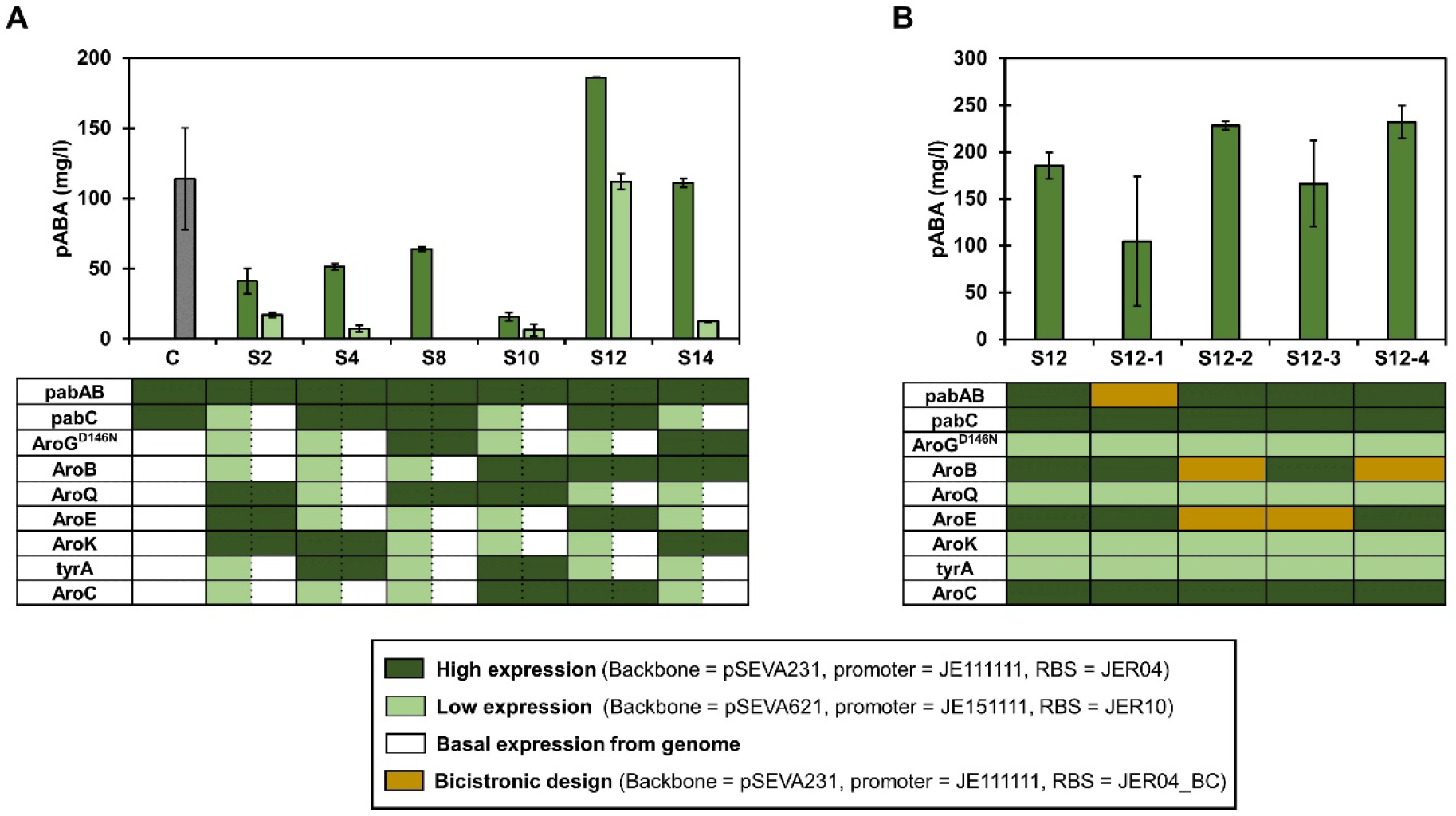
Expansion of the design space. **A**. Effect of reducing gene over-expression to native expression levels on pABA production. **B**. Effect of increasing expression levels using bicistronic designs on pABA production.

Bicistronic designs were used to study whether higher expression of *pabAB, aroB*, and *aroE*, the genes with the highest positive impact on pABA production, could further improve pABA titers. Four additional strains (S12-1, S12-2, S12-3, and S12-4) were constructed with bicistronic designs controlling the expression of *pabAB, aroB, aroE* or *aroB* and *aroE* using the best producer strain, S12, as background (Figure 3B). Higher expression of *aroB* in strain S12-4 resulted in a 25.2% increased production compared to S12 with a final titer of 232 mg/l and a yield on glucose of 0.024 mol/mol. These results prove *aroB* to be a limiting factor for pABA production in *P. putida*.

## 4 Discussion

Adjusting gene expression is one of the required steps to optimize chemical production in cell factories ^12,14,28^. However, in order to consider possible synergies among genes, numerous strains that vary in the over-expressed genes must be constructed. In this study, we used a Plackett-Burman design to screen for the effect of nine genes involved in pABA production. Considering only two over-expression levels per gene, testing all possible combinations of mild and high over-expression would require the construction of 512 strains. Instead, the DoE design proposes the construction of 16 strains (3% of all the combinations). Even though only 14 of the 16 strains could be constructed after repeated attempts, identifying the effect of each of the over-expressed genes on pABA production was possible by training a linear model on the production data, showcasing the robustness of the proposed design to missing data.

*P. putida* has been engineered for the production of many relevant aromatic-derived compounds. However, optimizing the shikimate pathway in this microorganism remains a challenge^2,17,29^. While over-expression of all the shikimate pathway genes, *pabAB* (heterodimer complex) and *pabC* is required to obtain higher pABA titers, we show that different overexpression strengths per gene are optimal. While mild overexpression of *tyrA, aroQ, aroG*^D146N^, and *aroC* is needed, higher over-expression of *pabABC, aroE*, and especially *aroB* is beneficial. The proposed approach can efficiently identify production bottlenecks caused by insufficient gene expression. Notably, although the expression of feed-back resistant variants of *aroG* is generally considered beneficial for production, high over-expression of this gene has a negative impact on pABA production in *P. putida*. In contrast, and as previously reported for muconic acid production in *P. putida*^30^ and violacein production in *E. coli*^5^, over-expression *aroB* is identified as limiting. The proposed approach resulted in a pABA production range from 2.6 ± 0.4 mg/l to 232.1 ± 17.6 mg/l among the 25 strains tested in minimal media with glucose. Furthermore, it enabled the identification of *aroB* expression, encoding 3-dehydroquinate synthase, as a significant limiting factor in the biosynthesis of this shikimate-derived product in *P. putida*. The reported titers are competitive compared to results obtained with other organisms such as yeast reaching 215 mg/l^18^. However, higher titers have been obtained in *E. coli* and *Corynebacterium glutamicum*, 4.8 g/l and 43 g/l, respectively, using rich media in fed-batch cultures^25,31^. Beyond the optimization of gene expression, *pabABC* gene has a significant impact on pABA production and more efficient heterologous genes could improve the titers obtained with *P. putida* ^25,31–33^. Besides, strategies such as improving precursor supply, optimizing production conditions, and deleting genes from competing pathways could further improve the strain performance ^17,32^. In addition, the use of fed-batch cultivations as part of the bioprocess optimization is an approach expected to yield higher product titers compared to the short batch cultivations in falcon tubes used in this study ^34^.

In conclusion, this study demonstrates an effective strategy to explore the metabolic pathway design for overproduction of pABA by manipulating the expression levels of *pabABC* and all shikimate pathway genes. We could identify *aroB* expression as a relevant limiting step in the accumulation of this aromatic compound in *P. putida*. Considering that combinatorial metabolic engineering is a requirement for strain optimization, we demonstrate the utility of this approach for the optimization of gene expression and the identification of bottlenecks that can serve as a starting point during cell factory design.

