## Supplementary material for "Combinatorial engineering reveals shikimate pathway bottlenecks in para-aminobenzoic acid production in *Pseudomonas putida*"

Supplementary materials

Table S1. List of strains used in this study

| **Strain** | **Characteristics and genotype** | **Reference** |
| --- | --- | --- |
| *Escherichia coli*  Dh5α | F-λ-*endA*1 *glnX*44(AS) *thiE*1 *recA*1 *relA*1 *spoT*1 *gyrA*96(NalR) *rfbC*1 *deoR* *nupG* Φ80(*lacZ*ΔM15) Δ(*argF*-*lac*)U169 *hsdR*17(rK mK ) | ^1^ |
| *Escherichia coli*  K-12 MG1655 | F- lambda- *ilvG*- *rfb*-50 *rph*-1 | ^2^ |
| *Pseudomonas*  *Putida* KT2440 | Wild-type strain; mt-2 derivative cured of the TOL plasmid pWW0 | ^3^ |

Table S2. List of plasmids used in this study

| **Name** | **Relevant features** | **Use** | **Reference** |
| --- | --- | --- | --- |
| pSEVAb231 | oriV(pBBR1) ; Km | High expression level | ^4^ |
| pSEVAb621 | oriV(RK2); Gm | moderate expression level | ^4^ |
| pSB1C3 | oriV(pMB1); CmR | *aroG*^D146N^ cloning | ^5^ |

Table S3. List of primers used in this study

| **Primer name** | **Sequence** (**5’→3’**) | **Template** | **Description and use** |
| --- | --- | --- | --- |
| backbone 1wgi fw | attcgcgtggcagaagagggcaagtcgtgatccggcaaaaaagggcaagg | pSEVAB621 | pSEVAb621 linearization |
| backbone 1wgi rv | atcatacctgacctccataggaaatcatccttagcgaaagctaa | pSEVAB621 | pSEVAb621 linearization |
| JE1511111 fw | ctttcgctaaggatgatttcctatggaggtcaggtatgatt | gBlock JE151111 | JE151111 linearization |
| JE1511111 rv | agttatctataagcaggatcatatgatactttcgtgtgtgagttaatcttaagaattgttatcc | gBlock JE151111 | JE151111 linearization |
| paba 1w fw | ttaagattaactcacacacgaaagtatcatatgatcctgcttatagataactacg | gDNA *E. coli* K-12 | pabA linearization |
| paba 1w rv | tcacagcgggagataacgtcttcatatgatactttcgttcagcgatgcaggaaattagcc | gDNA *E. coli* K-12 | pabA linearization |
| pabb 1w fw | atttcctgcatcgctgaacgaaagtatcatatgaagacgttatctcccgc | gDNA *E. coli* K-12 | pabB linearization |
| pabb 1w rv | atgatactttcgtttacttctccagttgcttc | gDNA *E. coli* K-12 | pabB linearization |
| pabc 1w fw | agcaactggagaagtaaacgaaagtatcatatgttcttaattaacggtcataagcag | gDNA *E. coli* K-12 | pabC linearization |
| pabc 1w rv | atgcgtaaatcgtcgttctgataattcatatgatactttcgtattcgggcgctcacaaag | gDNA *E. coli* K-12 | pabC linearization |
| arog 1w fw | tttgtgagcgcccgaatacgaaagtatcatatgaattatcagaacgacg | aroG^D146N^ plasmid | aroG^D146N^ linearization |
| arog 1w rv | atgatactttcgtttacccgcgacgcgcttttactgc | aroG^D146N^ plasmid | aroG^D146N^ linearization |
| arob 1wgib fw | aagcgcgtcgcgggtaaacgaaagtatcatatgcagacactaaaggtcgacctg | gDNA *P. putida* KT2440 | aroB linearization |
| arob 1wgib rv | atgatactttcgttcaaagctgggccacgatcgc | gDNA *P. putida* KT2440 | aroB linearization |
| aroq 1wgib fw | tcgtggcccagctttgaacgaaagtatcatatggcaacgctactggtgctcc | gDNA *P. putida* KT2440 | aroQ linearization |
| aroq 1wgib rv | atgatactttcgttcatttgggctgtgcgttggcagcc | gDNA *P. putida* KT2440 | aroQ linearization |
| aroe 1wgib fw | acgcacagcccaaatgaacgaaagtatcatatggaccagtacgtcgtttttgg | gDNA *P. putida* KT2440 | aroE linearization |
| aroe 1wgib rv | atgatactttcgttcagccccgagcgagttg | gDNA *P. putida* KT2440 | aroE linearization |
| arok 1wgib fw | aactcgctcggggctgaacgaaagtatcatgtgcgaaatttgatacttgtgg | gDNA *P. putida* KT2440 | aroK linearization |
| arok 1wgib rv | atgatactttcgtttaacggggcggcaactgctgc | gDNA *P. putida* KT2440 | aroK linearization |
| tyra 1wgib fw | agttgccgccccgttaaacgaaagtatcatgtggtaaatgcagcaaaaacc | gDNA *P. putida* KT2440 | tyrA linearization |
| tyra 1wgi rv | tgacaccttgcccttttttgccggatcacgacttgccctcttctgcc | gDNA *P. putida* KT2440 | tyrA linearization |
| backbone 1s fw | ccgggtctcaattccagaaatcatccttagcgaaagc | pSEVAB231 | pSEVAb231 linearization |
| backbone 1s rv | aggtctcacagtccggcaaaaaaggg | pSEVAB231 | pSEVAb231 linearization |
| JE1111111 fw | taggtctccactactatggaggtcaggtatg | gBlock JE111111 | JE111111 linearization |
| JE1111111 rv | taggtctcagggcgtgtgagttaatcttaagaattgttatccgc | gBlock JE111111 | JE111111 linearization |
| aroc 1s fw | taggtctcagcccgacgagggtatcatatgtccggcaatacctacgg | gDNA *P. putida* KT2440 | aroC linearization |
| aroc 1s rv | aggtctcgactgcagcggccgctactagtattattatcagcgctggcccagcaccggggtattc | gDNA *P. putida* KT2440 | aroC linearization |
| pabc 2w fw | taggtctcagcccgacgaaagtatcatatgttcttaattaacggtcataagcagg | gDNA *E. coli* K-12 | pabC linearization |
| pabc 2w rv | taggtctcataccattcgggcgctcacaaag | gDNA *E. coli* K-12 | pabC linearization |
| arog 2w fw | taggtctcaggtagacgaaagtatcatatgaattatcagaacgacg | aroG^D146N^ plasmid | aroG^D146N^ linearization |
| arog 2w rv | taggtctcaccgtttacccgcgacgcgcttttactgc | aroG^D146N^ plasmid | aroG^D146N^ linearization |
| arob 2w fw | taGGTCTCaACGGgACGAAAGtatcatatgcagacactaaaggtcgacctg | gDNA *P. putida* KT2440 | aroB linearization |
| arob 2w rv | taggtctcaaccgtcaaagctgggccacgatcgc | gDNA *P. putida* KT2440 | aroB linearization |
| tyra 2w fw | taggtctcacggtgacgagggtatcatgtggtaaatgcagcaaaaacc | gDNA *P. putida* KT2440 | tyrA linearization |
| tyra 2w rv | taggtctcaagtatcacgacttgccctcttctgcc | gDNA *P. putida* KT2440 | tyrA linearization |
| aroc 2w fw | taggtctcatactgacgaaagtatcatatgtccggcaatacctacgg | gDNA *P. putida* KT2440 | aroC linearization |
| aroc 2w rv | aggtctcgactgcagcggccgctactagtattattatcagcgctggcccagcaccggggtattc | gDNA *P. putida* KT2440 | aroC linearization |
| paba 2s fw | taggtctcagcccgacgagggtatcatatgatcctgcttatagataactacg | gDNA *E. coli* K-12 | pabA linearization |
| paba 2s rv | gtggtctctttggtcagcgatgcaggaaattagcc | gDNA *E. coli* K-12 | pabA linearization |
| pabb 2s fw | taggtctcaccaagacgagggtatcatatgaagacgttatctcccgc | gDNA *E. coli* K-12 | pabB linearization |
| pabb 2s rv | gtggtctcttctgttacttctccagttgcttc | gDNA *E. coli* K-12 | pabB linearization |
| aroq 2s fw | taggtctcacagagacgagggtatcatatggcaacgctactggtgctcc | gDNA *P. putida* KT2440 | aroQ linearization |
| aroq 2s rv | taggtctcagttctcatttgggctgtgcgttggcagc | gDNA *P. putida* KT2440 | aroQ linearization |
| aroe 2s fw | taggtctcagaacgacgagggtatcatatggaccagtacgtcgtttttgg | gDNA *P. putida* KT2440 | aroE linearization |
| aroe 2s rv | taggtctcatgactcagccccgagcgagttg | gDNA *P. putida* KT2440 | aroE linearization |
| arok 2s fw | taggtctcagtcagacgagggtatcatgtgcgaaatttgatacttgtgg | gDNA *P. putida* KT2440 | aroK linearization |
| arok 2s rv | taggtctcaagtattaacggggcggcaactgctgc | gDNA *P. putida* KT2440 | aroK linearization |
| paba 3w fw | taggtctcagcccgacgaaagtatcatatgatcctgcttatagataactacg | gDNA E. coli K-12 | pabA linearization |
| paba 3w rv | gtggtctctttggtcagcgatgcaggaaattagcc | gDNA *E. coli* K-12 | pabA linearization |
| pabb 3w fw | taggtctcaccaagacgaaagtatcatatgaagacgttatctcccgc | gDNA *E. coli* K-12 | pabB linearization |
| pabb 3w rv | gtggtctctggatttacttctccagttgcttc | gDNA *E. coli* K-12 | pabB linearization |
| arog 3w fw | taggtctcaatccgacgaaagtatcatatgaattatcagaacgacg | aroG^D146N^ plasmid | aroG^D146N^ linearization |
| arog 3w rv | taggtctcaccgtttacccgcgacgcgcttttactgc | aroG^D146N^ plasmid | aroG^D146N^ linearization |
| arob 3w fw | taggtctcaacgggacgaaagtatcatatgcagacactaaaggtcgacctg | gDNA *P. putida* KT2440 | aroB linearization |
| arob 3w rv | taggtctcaaccgtcaaagctgggccacgatcgc | gDNA *P. putida* KT2440 | aroB linearization |
| arok 3w fw | taggtctcacggtgacgaaagtatcatgtgcgaaatttgatacttgtgg | gDNA *P. putida* KT2440 | aroK linearization |
| arok 3w rv | taggtctcaagtattaacggggcggcaactgctgc | gDNA *P. putida* KT2440 | aroK linearization |
| aroc 3w fw | taggtctcatactgacgaaagtatcatatgtccggcaatacctacgg | gDNA *P. putida* KT2440 | aroC linearization |
| aroc pp 3w rv | aggtctcgactgcagcggccgctactagtattattatcagcgctggcccagcaccggggtattc | gDNA *P. putida* KT2440 | aroC linearization |
| pabc 3s fw | taggtctcagcccgacgagggtatcatatgttcttaattaacggtcataagcagg | gDNA *E. coli* K-12 | pabC linearization |
| pabc 3s rv | taggtctcatctgattcgggcgctcacaaag | gDNA *E. coli* K-12 | pabC linearization |
| aroq 3s fw | taggtctcacagagacgagggtatcatatggcaacgctactggtgc | gDNA *P. putida* KT2440 | aroQ linearization |
| aroq 3s rv | taggtctcagttctcatttgggctgtgcgttgg | gDNA *P. putida* KT2440 | aroQ linearization |
| aroe 3s fw | taggtctcagaacgacgagggtatcatatggaccagtacgtcgtttttgg | gDNA *P. putida* KT2440 | aroE linearization |
| aroe 3s rv | taggtctcatgactcagccccgagcgagttg | gDNA *P. putida* KT2440 | aroE linearization |
| tyra 3s fw | aactcgctcggggctgaacgagggtatcatgtggtaaatgcagcaaaaacc | gDNA *P. putida KT2440* | tyrA linearization |
| tyra 3s rv | taggtctcaagtatcacgacttgccctcttctgcc | gDNA *P. putida* KT2440 | tyrA linearization |
| arog 4wgib fw | ttaagattaactcacacacgaaagtatcatatgaattatcagaacgacg | aroG^D146N^ plasmid | aroG^D146N^ linearization |
| arog 4wgib rv | atgatactttcgtttacccgcgacgcgcttttactg | aroG^D146N^ plasmid | aroG^D146N^ linearization |
| arob 4wgib fw | aagcgcgtcgcgggtaaacgaaagtatcatatgcagacactaaaggtcg | gDNA *P. putida* KT2440 | aroB linearization |
| arob 4wgib rv | atgatactttcgttcaaagctgggccacgatcg | gDNA *P. putida* KT2440 | aroB linearization |
| aroq 4wgib fw | tcgtggcccagctttgaacgaaagtatcatatggcaacgctactggtgc | gDNA *P. putida* KT2440 | aroQ linearization |
| aroq 4wgib rv | atgatactttcgttcatttgggctgtgcgttgg | gDNA *P. putida* KT2440 | aroQ linearization |
| aroe 4wgib fw | acgcacagcccaaatgaacgaaagtatcatatggaccagtacgtcgtttttgg | gDNA *P. putida* KT2440 | aroE linearization |
| aroe 4wgib rv | atgatactttcgttcagccccgagcgagttg | gDNA *P. putida* KT2440 | aroE linearization |
| aroc 4wgib fw | aactcgctcggggctgaacgaaagtatcatatgtccggcaatacctacg | gDNA *P. putida* KT2440 | aroC linearization |
| aroc 4wgib rv |  | gDNA *P. putida* KT2440 | aroC linearization |
| paba 4sgib fw | ttaagattaactcacacacgagggtatcatatgatcctgcttatagataactacg | gDNA *E. coli* K-12 | pabA linearization |
| paba 4sgib rv | atgataccctcgttcagcgatgcaggaaattagcc | gDNA *E. coli* K-12 | pabA linearization |
| pabb 4sgib fw | atttcctgcatcgctgaacgagggtatcatatgaagacgttatctcccgc | gDNA *E. coli* K-12 | pabB linearization |
| pabb 4sgib rv | atgataccctcgtttacttctccagttgcttc | gDNA *E. coli* K-12 | pabB linearization |
| pabc 4sgib fw | agcaactggagaagtaaacgagggtatcatatgttcttaattaacggtcataagcag | gDNA *E. coli* K-12 | pabC linearization |
| pabc 4sgib rv | atgataccctcgtattcgggcgctcacaaag | gDNA *E. coli* K-12 | pabC linearization |
| arok 4sgib fw | tttgtgagcgcccgaatacgagggtatcatgtgcgaaatttgatacttgtgg | gDNA *P. putida* KT2440 | aroK linearization |
| arok 4sgib rv | atgataccctcgtttaacggggcggcaactgctgc | gDNA *P. putida* KT2440 | aroK linearization |
| tyra 4sgib fw | agttgccgccccgttaaacgagggtatcatgtggtaaatgcagcaaaaacc | gDNA *P. putida* KT2440 | tyrA linearization |
| tyra 4sgib rv | ccttgcccttttttgccggatcacgacttgccctcttctgcc | gDNA *P. putida* KT2440 | tyrA linearization |
| paba 5wgib fw | ttaagattaactcacacacgaaagtatcatatgatcctgcttatagataactac | gDNA *E. coli* K-12 | pabA linearization |
| paba 5wgib rv | atgatactttcgttcagcgatgcaggaaattagcc | gDNA *E. coli* K-12 | pabA linearization |
| pabb 5wgib fw | atttcctgcatcgctgaacgaaagtatcatatgaagacgttatctcccgc | gDNA *E. coli* K-12 | pabB linearization |
| pabb 5wgib rv | atgatactttcgtttacttctccagttgcttc | gDNA *E. coli* K-12 | pabB linearization |
| pabc 5wgib fw | agcaactggagaagtaaacgaaagtatcatatgttcttaattaacggtcataagcag | gDNA *E. coli* K-12 | pabC linearization |
| pabc 5wgib rv | atgatactttcgtattcgggcgctcacaaag | gDNA *E. coli* K-12 | pabC linearization |
| arob 5wgib fw | tttgtgagcgcccgaatacgaaagtatcatatgcagacactaaaggtcgacctg | gDNA *P. putida* KT2440 | aroB linearization |
| arob 5wgib rv | atgatactttcgttcaaagctgggccacgatcgc | gDNA *P. putida* KT2440 | aroB linearization |
| aroe 5wgib fw | tcgtggcccagctttgaacgaaagtatcatatggaccagtacgtcgtttttgg | gDNA *P. putida* KT2440 | aroE linearization |
| aroe 5w gib rv | ccttgcccttttttgccggatcagccccgagcgagttg | gDNA *P. putida* KT2440 | aroE linearization |
| arog 5sgib fw | ttaagattaactcacacacgagggtatcatatgaattatcagaacgacg | aroG^D146N^ plasmid | aroG^D146N^ linearization |
| arog 5sgib rv | atgataccctcgtttacccgcgacgcgcttttactgc | aroG^D146N^ plasmid | aroG^D146N^ linearization |
| aroq 5sgib fw | aagcgcgtcgcgggtaaacgagggtatcatatggcaacgctactggtgctcc | gDNA *P. putida* KT2440 | aroQ linearization |
| aroq 5sgib rv | atgataccctcgttcatttgggctgtgcgttggcagcc | gDNA *P. putida* KT2440 | aroQ linearization |
| arok 5sgib fw | acgcacagcccaaatgaacgagggtatcatgtgcgaaatttgatacttgtgg | gDNA *P. putida* KT2440 | aroK linearization |
| arok 5sgib rv | atgataccctcgtttaacggggcggcaactgctgc | gDNA *P. putida* KT2440 | aroK linearization |
| tyra 5sgib fw | agttgccgccccgttaaacgagggtatcatgtggtaaatgcagcaaaaacc | gDNA *P. putida* KT2440 | tyrA linearization |
| tyra 5sgib rv | atgataccctcgttcacgacttgccctcttctgcc | gDNA *P. putida* KT2440 | tyrA linearization |
| aroc 5sgib fw | aagagggcaagtcgtgaacgagggtatcatatgtccggcaatacctacggcaagc | gDNA *P. putida* KT2440 | aroC linearization |
| aroc 5sgib rv | ccttgcccttttttgccggatcagcgctggcccagcaccg | gDNA *P. putida* KT2440 | aroC linearization |
| pabc 6wgib fw | ttaagattaactcacacacgaaagtatcatatgttcttaattaacggtcataagc | gDNA *E. coli* K-12 | pabC linearization |
| pabc 6wgib rv | atgatactttcgtattcgggcgctcacaaagtg | gDNA *E. coli* K-12 | pabC linearization |
| arob 6wgib fw | tttgtgagcgcccgaatACGAAAGtatcatatgcagacactaaaggtcg | gDNA *P. putida* KT2440 | aroB linearization |
| arob 6wgib rv | atgatactttcgttcaaagctgggccacgatcg | gDNA *P. putida* KT2440 | aroB linearization |
| aroq 6wgib fw | tcgtggcccagctttgaacgaaagtatcatatggcaacgctactggtgc | gDNA *P. putida* KT2440 | aroQ linearization |
| aroq 6wgib rv | atgatactttcgttcatttgggctgtgcgttg | gDNA *P. putida* KT2440 | aroQ linearization |
| arok 6wgib fw | acgcacagcccaaatgaacgaaagtatcatgtgcgaaatttgatacttgtgg | gDNA *P. putida* KT2440 | aroK linearization |
| arok 6w gib rv | ccttgcccttttttgccggattaacggggcggcaactgc | gDNA *P. putida* KT2440 | aroK linearization |
| paba 6sgib fw | ttaagattaactcacacacgagggtatcatatgatcctgcttatagataactacg | gDNA *E. coli* K-12 | pabA linearization |
| paba 6sgib rv | atgataccctcgttcagcgatgcaggaaattagcc | gDNA *E. coli* K-12 | pabA linearization |
| pabb 6sgib fw | atttcctgcatcgctgaacgagggtatcatatgaagacgttatctcccgc | gDNA *E. coli* K-12 | pabB linearization |
| pabb 6sgib rv | atgataccctcgtttacttctccagttgcttc | gDNA *E. coli* K-12 | pabB linearization |
| arog 6sgib fw | agcaactggagaagtaaacgagggtatcatatgaattatcagaacgacg | aroG^D146N^ plasmid | aroG^D146N^ linearization |
| arog 6sgib rv | atgataccctcgtttacccgcgacgcgcttttactgc | aroG^D146N^ plasmid | aroG^D146N^ linearization |
| aroe 6sgib fw | aagcgcgtcgcgggtaaacgagggtatcatatggaccagtacgtcgtttttgg | gDNA *P. putida* KT2440 | aroE linearization |
| aroe 6sgib rv | atgataccctcgttcagccccgagcgagttg | gDNA *P. putida* KT2440 | aroE linearization |
| tyra 6sgib fw | aactcgctcggggctgaacgagggtatcatgtggtaaatgcagcaaaaacc | gDNA *P. putida* KT2440 | tyrA linearization |
| tyra 6sgib rv | atgataccctcgttcacgacttgccctcttctgcc | gDNA *P. putida* KT2440 | tyrA linearization |
| aroc 6sgib fw | aagagggcaagtcgtgaacgagggtatcatatgtccggcaatacctacggcaagc | gDNA P. putida KT2440 | aroC linearization |
| aroc 6sgbi rv | tgcagcggccgctactagtatcagcgctggcccagcaccg | gDNA P. putida KT2440 | aroC linearization |
| paba 7wgib fw | ttaagattaactcacacacgaaagtatcatatgatcctgcttatagataactacg | gDNA *E. coli* K-12 | pabA linearization |
| paba 7wgib rv | atgatactttcgttcagcgatgcaggaaattagcc | gDNA *E. coli* K-12 | pabA linearization |
| pabb 7wgib fw | atttcctgcatcgctgaacgaaagtatcatatgaagacgttatctcccgc | gDNA *E. coli* K-12 | pabB linearization |
| pabb 7wgib rv | atgatactttcgtttacttctccagttgcttc | gDNA *E. coli* K-12 | pabB linearization |
| arob 7wgib fw | agcaactggagaagtaaacgaaagtatcatatgcagacactaaaggtcgacctg | gDNA *P. putida* KT2440 | aroB linearization |
| arob 7wgib rv | atgatactttcgttcaaagctgggccacgatcgc | gDNA *P. putida* KT2440 | aroB linearization |
| aroq 7wgib fw | tcgtggcccagctttgaacgaaagtatcatatggcaacgctactggtgctcc | gDNA *P. putida* KT2440 | aroQ linearization |
| aroq 7wgib rv | atgatactttcgttcatttgggctgtgcgttggcagcc | gDNA *P. putida* KT2440 | aroQ linearization |
| tyra 7wgib fw | acgcacagcccaaatgaacgaaagtatcatgtggtaaatgcagcaaaaacc | gDNA *P. putida* KT2440 | tyrA linearization |
| tyra 7wgi rv | tgacaccttgcccttttttgccggatcacgacttgccctcttctgcc | gDNA *P. putida* KT2440 | tyrA linearization |
| pabc 7s fw | taggtctcagcccgacgagggtatcatatgttcttaattaacggtcataagcagg | gDNA *E. coli* K-12 | pabC linearization |
| pabc 7s rv | taggtctcatctgattcgggcgctcacaaag | gDNA *E. coli* K-12 | pabC linearization |
| arog 7s fw | taggtctcacagagacgagggtatcatatgaattatcagaacgacg | aroG^D146N^ plasmid | aroG^D146N^ linearization |
| arog 7s rv | taggtctcagttcttacccgcgacgcgcttttactgc | aroG^D146N^ plasmid | aroG^D146N^ linearization |
| aroe 7s fw | taggtctcagaacgacgagggtatcatatggaccagtacgtcgtttttgg | gDNA *P. putida* KT2440 | aroE linearization |
| aroe 7s rv | taggtctcatgactcagccccgagcgagttg | gDNA *P. putida* KT2440 | aroE linearization |
| arok 7s fw | taggtctcagtcagacgagggtatcatgtgcgaaatttgatacttgtgg | gDNA *P. putida* KT2440 | aroK linearization |
| arok 7s rv | taggtctcaagtattaacggggcggcaactgctgc | gDNA *P. putida* KT2440 | aroK linearization |
| aroc 7s fw | taggtctcatactgacgagggtatcatatgtccggcaatacctacgg | gDNA *P. putida* KT2440 | aroC linearization |
| aroc 7s rv | aggtctcgactgcagcggccgctactagtattattatcagcgctggcccagcaccggggtattc | gDNA *P. putida* KT2440 | aroC linearization |
| arob 8wgib fw | ttaagattaactcacacacgaaagtatcatatgcagacactaaaggtcgacctg | gDNA *P. putida* KT2440 | aroB linearization |
| arob 8wgib rv | atgatactttcgttcaaagctgggccacgatcgc | gDNA *P. putida* KT2440 | aroB linearization |
| aroe 8wgib fw | tcgtggcccagctttgaacgaaagtatcatatggaccagtacgtcgtttttgg | gDNA *P. putida* KT2440 | aroE linearization |
| aroe 8wgib rv | atgatactttcgttcagccccgagcgagttg | gDNA *P. putida* KT2440 | aroE linearization |
| arok 8wgib fw | aactcgctcggggctgaacgaaagtatcatgtgcgaaatttgatacttgtgg | gDNA *P. putida* KT2440 | aroK linearization |
| arok 8wgib rv | atgatactttcgtttaacggggcggcaactgctgc | gDNA *P. putida* KT2440 | aroK linearization |
| tyra 8wgib fw | agttgccgccccgttaaacgaaagtatcatgtggtaaatgcagcaaaaacc | gDNA *P. putida* KT2440 | tyrA linearization |
| tyra 8wgib rv | atgatactttcgttcacgacttgccctcttctgcc | gDNA *P. putida* KT2440 | tyrA linearization |
| aroc 8wgib fw | aagagggcaagtcgtgaacgaaagtatcatatgtccggcaatacctacggcaagc | gDNA *P. putida* KT2440 | aroC linearization |
| aroc 8wgi rv | ccttgcccttttttgccggatcagcgctggcccagcaccg | gDNA *P. putida* KT2440 | aroC linearization |
| paba 8sgib fw | ttaagattaactcacacacgagggtatcatatgatcctgcttatagataactacg | gDNA *E. coli* K-12 | pabA linearization |
| paba 8sgib rv | atgataccctcgttcagcgatgcaggaaattagcc | gDNA *E. coli* K-12 | pabA linearization |
| pabb 8sgib fw | atttcctgcatcgctgaacgagggtatcatatgaagacgttatctcccgc | gDNA *E. coli* K-12 | pabB linearization |
| pabb 8sgib rv | atgataccctcgtttacttctccagttgcttc | gDNA *E. coli* K-12 | pabB linearization |
| pabc 8sgib fw | agcaactggagaagtaaacgagggtatcatatgttcttaattaacggtcataagcag | gDNA *E. coli* K-12 | pabC linearization |
| pabc 8sgib rv | atgataccctcgtattcgggcgctcacaaag | gDNA E. coli K-12 | pabC linearization |
| arog 8sgib fw | ttagccccactttgtgagcgcccgaatacgagggtatcatatgaattatcagaacgacg | aroG^D146N^ plasmid | aroG^D146N^ linearization |
| arog 8sgib rv | atgataccctcgtttacccgcgacgcgcttttactgc | aroG^D146N^ plasmid | aroG^D146N^ linearization |
| aroq 8sgib fw | aagcgcgtcgcgggtaaacgagggtatcatatggcaacgctactggtgctcc | gDNA *P. putida* KT2440 | aroQ linearization |
| aroq 8sgib rv | atgataccctcgttcatttgggctgtgcgttggcagcc | gDNA *P. putida* KT2440 | aroQ linearization |
| paba 9wgi fw | aattcttaagattaactcacacacgaaagtatcatatgatcctgcttatagataactacg | gDNA *E. coli* K-12 | pabA linearization |
| paba 9wgib rv | atgatactttcgttcagcgatgcaggaaattagcc | gDNA *E. coli* K-12 | pabA linearization |
| pabb 9wgib fw | atttcctgcatcgctgaacgaaagtatcatatgaagacgttatctcccgc | gDNA *E. coli* K-12 | pabB linearization |
| pabb 9wgib rv | atgatactttcgtttacttctccagttgcttc | gDNA *E. coli* K-12 | pabB linearization |
| pabc 9wgib fw | agcaactggagaagtaaacgaaagtatcatatgttcttaattaacggtcataagcag | gDNA *E. coli* K-12 | pabC linearization |
| pabc 9wgib rv | atgatactttcgtattcgggcgctcacaaag | gDNA *E. coli* K-12 | pabC linearization |
| arog 9wgib fw | tttgtgagcgcccgaatacgaaagtatcatatgaattatcagaacgacg | aroG^D146N^ plasmid | aroG^D146N^ linearization |
| arog 9wgib rv | atgatactttcgtttacccgcgacgcgcttttactgc | aroG^D146N^ plasmid | aroG^D146N^ linearization |
| aroq 9wgib fw | aagcgcgtcgcgggtaaacgaaagtatcatatggcaacgctactggtgctcc | gDNA *P. putida* KT2440 | aroQ linearization |
| aroq 9wgib rv | atgatactttcgttcatttgggctgtgcgttggcagcc | gDNA *P. putida* KT2440 | aroQ linearization |
| aroc 9wgib fw | acgcacagcccaaatgaacgaaagtatcatatgtccggcaatacctacggcaagc | gDNA *P. putida* KT2440 | aroC linearization |
| aroc 9wgi rv | ccttgcccttttttgccggatcagcgctggcccagcaccg | gDNA *P. putida* KT2440 | aroC linearization |
| arob 9sgib fw | ttaagattaactcacacacgagggtatcatatgcagacactaaaggtcgacctg | gDNA *P. putida* KT2440 | aroB linearization |
| arob 9sgib rv | atgataccctcgttcaaagctgggccacgatcgc | gDNA *P. putida* KT2440 | aroB linearization |
| aroe 9sgib fw | tcgtggcccagctttgaacgagggtatcatatggaccagtacgtcgtttttgg | gDNA *P. putida* KT2440 | aroE linearization |
| aroe 9sgib rv | atgataccctcgttcagccccgagcgagttg | gDNA *P. putida* KT2440 | aroE linearization |
| arok 9sgib fw | aactcgctcggggctgaacgagggtatcatgtgcgaaatttgatacttgtgg | gDNA *P. putida* KT2440 | aroK linearization |
| arok 9sgib rv | atgataccctcgtttaacggggcggcaactgctgc | gDNA *P. putida* KT2440 | aroK linearization |
| tyra 9sgib fw | agttgccgccccgttaaacgagggtatcatgtggtaaatgcagcaaaaacc | gDNA *P. putida* KT2440 | tyrA linearization |
| tyra 9sgib rv | ccttgcccttttttgccggatcacgacttgccctcttctg | gDNA *P. putida* KT2440 | tyrA linearization |
| pabc 10wgib fw | ttaagattaactcacacacgaaagtatcatatgttcttaattaacggtcataagcag | gDNA *E. coli* K-12 | pabC linearization |
| pabc 10wgib fw | agcaactggagaagtaaacgaaagtatcatatgttcttaattaacggtcataagcag | gDNA *E. coli* K-12 | pabC linearization |
| arog 10wgib fw | tttgtgagcgcccgaatacgaaagtatcatatgaattatcagaacgacg | aroG^D146N^ plasmid | aroG^D146N^ linearization |
| arog 10wgib rv | atgatactttcgtttacccgcgacgcgcttttactgc | aroG^D146N^ plasmid | aroG^D146N^ linearization |
| aroe 10wgib fw | aagcgcgtcgcgggtaaacgaaagtatcatatggaccagtacgtcgtttttgg | gDNA *P. putida* KT2440 | aroE linearization |
| aroe 10wgib rv | atgatactttcgttcagccccgagcgagttg | gDNA *P. putida* KT2440 | aroE linearization |
| arok 10wgib fw | aactcgctcggggctgaacgaaagtatcatgtgcgaaatttgatacttgtgg | gDNA *P. putida* KT2440 | aroK linearization |
| arok 10wgi rv | gtttggtttttgctgcatttaccacatgatactttcgtttaacggggcggcaactgctgc | gDNA *P. putida* KT2440 | aroK linearization |
| paba 10sgib fw | ttaagattaactcacacacgagggtatcatatgatcctgcttatagataactacg | gDNA *E. coli* K-12 | pabA linearization |
| paba 10sgib rv | atgataccctcgttcagcgatgcaggaaattagcc | gDNA *E. coli* K-12 | pabA linearization |
| pabb 10sgib fw | atttcctgcatcgctgaacgagggtatcatatgaagacgttatctcccgc | gDNA *E. coli* K-12 | pabB linearization |
| pabb 10sgib rv | atgataccctcgtttacttctccagttgcttc | gDNA *E. coli* K-12 | pabB linearization |
| arob 10sgib fw | agcaactggagaagtaaacgagggtatcatatgcagacactaaaggtcgacctg | gDNA *P. putida* KT2440 | aroB linearization |
| arob 10sgib rv | atgataccctcgttcaaagctgggccacgatcgc | gDNA *P. putida* KT2440 | aroB linearization |
| aroq 10sgib fw | tcgtggcccagctttgaacgagggtatcatatggcaacgctactggtgctcc | gDNA *P. putida* KT2440 | aroQ linearization |
| aroq 10sgib rv | atgataccctcgttcatttgggctgtgcgttggcagcc | gDNA *P. putida* KT2440 | aroQ linearization |
| tyra 10sgib fw | acgcacagcccaaatgaacgagggtatcatgtggtaaatgcagcaaaaacc | gDNA *P. putida* KT2440 | tyrA linearization |
| tyra 10sgib rv | atgataccctcgttcacgacttgccctcttctgcc | gDNA *P. putida* KT2440 | tyrA linearization |
| aroc 10sgib fw | aagagggcaagtcgtgaacgagggtatcatatgtccggcaatacctacggcaagc | gDNA *P. putida* KT2440 | aroC linearization |
| aroc pp 10s rv | aggtctcgactgcagcggccgctactagtattattatcagcgctggcccagcaccggggtattc | gDNA *P. putida* KT2440 | aroC linearization |
| paba 11w fw | taggtctcagcccgacgaaagtatcatatgatcctgcttatagataactacg | gDNA *E. coli* K-12 | pabA linearization |
| paba 11w rv | gtggtctctttggtcagcgatgcaggaaattagcc | gDNA *E. coli* K-12 | pabA linearization |
| pabb 11w fw | taggtctcaccaagacgaaagtatcatatgaagacgttatctcccgc | gDNA *E. coli* K-12 | pabB linearization |
| pabb 11w rv | gtggtctcttaccttacttctccagttgcttc | gDNA *E. coli* K-12 | pabB linearization |
| arog 11w fw | taggtctcaggtagacgaaagtatcatatgaattatcagaacgacg | aroG^D146N^ plasmid | aroG^D146N^ linearization |
| arog 11w rv | aaaacgacgtactggtccatatgatactttcgtttacccgcgacgcgcttttactgc | aroG^D146N^ plasmid | aroG^D146N^ linearization |
| aroe 11w fw | taggtctcacggtgacgaaagtatcatatggaccagtacgtcgtttttgg | gDNA *P. putida* KT2440 | aroE linearization |
| aroe 11w rv | taggtctcatgactcagccccgagcgagttg | gDNA *P. putida* KT2440 | aroE linearization |
| tyra 11w fw | taggtctcagtcagacgaaagtatcatgtggtaaatgcagcaaaaacc | gDNA *P. putida* KT2440 | tyrA linearization |
| tyra 11w rv | taggtctcaagtatcacgacttgccctcttctgcc | gDNA *P. putida* KT2440 | tyrA linearization |
| pabc 11s fw | taggtctcagcccgacgagggtatcatatgttcttaattaacggtcataagcagg | gDNA E. coli K-12 | pabC linearization |
| pabc 11s rv | taggtctcatctgattcgggcgctcacaaag | gDNA E. coli K-12 | pabC linearization |
| arob 11s fw | taggtctcacagagacgagggtatcatatgcagacactaaaggtcgacctg | gDNA *P. putida* KT2440 | aroB linearization |
| arob 11s rv | taggtctcaccgttcaaagctgggccacgatcgc | gDNA *P. putida* KT2440 | aroB linearization |
| aroq 11s fw | taggtctcaacgggacgagggtatcatatggcaacgctactggtgctcc | gDNA *P. putida* KT2440 | aroQ linearization |
| aroq 11s rv | taggtctcatgactcatttgggctgtgcgttggcagc | gDNA *P. putida* KT2440 | aroQ linearization |
| arok 11s fw | taggtctcagtcagacgagggtatcatgtgcgaaatttgatacttgtgg | gDNA *P. putida* KT2440 | aroK linearization |
| arok 11s rv | taggtctcaagtattaacggggcggcaactgctgc | gDNA *P. putida* KT2440 | aroK linearization |
| aroc 11s fw | taggtctcatactgacgagggtatcatatgtccggcaatacctacgg | gDNA *P. putida* KT2440 | aroC linearization |
| aroc 11s rv | aggtctcgactgcagcggccgctactagtattattatcagcgctggcccagcaccggggtattc | gDNA *P. putida* KT2440 | aroC linearization |
| arog 12wgib fw | ttaagattaactcacacacgaaagtatcatatgaattatcagaacgacg | aroG^D146N^ plasmid | aroG^D146N^ linearization |
| arog 12wgib rv | atgatactttcgtttacccgcgacgcgcttttactgc | aroG^D146N^ plasmid | aroG^D146N^ linearization |
| aroq 12wgib fw | aagcgcgtcgcgggtaaacgaaagtatcatatggcaacgctactggtgctcc | gDNA *P. putida* KT2440 | aroQ linearization |
| aroq 12wgib rv | atgatactttcgttcatttgggctgtgcgttggcagcc | gDNA *P. putida* KT2440 | aroQ linearization |
| arok 12wgib fw | acgcacagcccaaatgaacgaaagtatcatgtgcgaaatttgatacttgtgg | gDNA *P. putida* KT2440 | aroK linearization |
| arok 12wgib rv | atgatactttcgtttaacggggcggcaactgctgc | gDNA *P. putida* KT2440 | aroK linearization |
| tyra 12wgib fw | agttgccgccccgttaaacgaaagtatcatgtggtaaatgcagcaaaaacc | gDNA *P. putida* KT2440 | tyrA linearization |
| tyra 12wgi rv | ccttgcccttttttgccggatcacgacttgccctcttctg | gDNA *P. putida* KT2440 | tyrA linearization |
| paba 12sgib fw | ttaagattaactcacacacgagggtatcatatgatcctgcttatagataactacg | gDNA *E. coli* K-12 | pabA linearization |
| paba 12sgib rv | atgataccctcgttcagcgatgcaggaaattagcc | gDNA *E. coli* K-12 | pabA linearization |
| pabb 12sgib fw | atttcctgcatcgctgaacgagggtatcatatgaagacgttatctcccgc | gDNA *E. coli* K-12 | pabB linearization |
| pabb 12sgib rv | atgataccctcgtttacttctccagttgcttc | gDNA *E. coli* K-12 | pabB linearization |
| pabc 12sgib fw | agcaactggagaagtaaacgagggtatcatatgttcttaattaacggtcataagcag | gDNA *E. coli* K-12 | pabC linearization |
| pabc 12sgib rv | atgataccctcgtattcgggcgctcacaaag | gDNA *E. coli* K-12 | pabC linearization |
| arob 12sgib fw | tttgtgagcgcccgaatacgagggtatcatatgcagacactaaaggtcgacctg | gDNA *P. putida* KT2440 | aroB linearization |
| arob 12sgib rv | atgataccctcgttcaaagctgggccacgatcgc | gDNA *P. putida* KT2440 | aroB linearization |
| aroe 12sgib fw | tcgtggcccagctttgaacgagggtatcatatggaccagtacgtcgtttttgg | gDNA *P. putida* KT2440 | aroE linearization |
| aroe 12sgib rv | atgataccctcgttcagccccgagcgagttg | gDNA *P. putida* KT2440 | aroE linearization |
| aroc 12sgib fw | gctcggggctgaacgagggtatcatatgtccggcaatacctacggcaagc | gDNA *P. putida* KT2440 | aroC linearization |
| aroc 12sgib rv | ccttgcccttttttgccggatcagcgctggcccagcaccg | gDNA *P. putida* KT2440 | aroC linearization |
| paba 13wgib fw | ttaagattaactcacacacgaaagtatcatatgatcctgcttatagataactacg | gDNA *E. coli* K-12 | pabA linearization |
| paba 13wgib rv | atgatactttcgttcagcgatgcaggaaattagcc | gDNA *E. coli* K-12 | pabA linearization |
| pabb 13wgib fw | atttcctgcatcgctgaacgaaagtatcatatgaagacgttatctcccgc | gDNA *E. coli* K-12 | pabB linearization |
| pabb 13wgib rv | atgatactttcgtttacttctccagttgcttc | gDNA *E. coli* K-12 | pabB linearization |
| pabc 13wgib fw | agcaactggagaagtaaacgaaagtatcatatgttcttaattaacggtcataagcag | gDNA *E. coli* K-12 | pabC linearization |
| pabc 13wgib rv | atgatactttcgtattcgggcgctcacaaag | gDNA *E. coli* K-12 | pabC linearization |
| arok 13wgib fw | tttgtgagcgcccgaatacgaaagtatcatgtgcgaaatttgatacttgtgg | gDNA *P. putida* KT2440 | aroK linearization |
| arok 13wgib rv | atgatactttcgtttaacggggcggcaactgctgc | gDNA *P. putida* KT2440 | aroK linearization |
| tyra 13wgib fw | agttgccgccccgttaaacgaaagtatcatgtggtaaatgcagcaaaaacc | gDNA *P. putida* KT2440 | tyrA linearization |
| tyra 13wgib rv | atgatactttcgttcacgacttgccctcttctgcc | gDNA *P. putida* KT2440 | tyrA linearization |
| aroc 13wgib fw | aagagggcaagtcgtgaacgaaagtatcatatgtccggcaatacctacggcaagc | gDNA *P. putida* KT2440 | aroC linearization |
| aroc 13wgi rv | ccttgcccttttttgccggatcagcgctggcccagcaccg | gDNA *P. putida* KT2440 | aroC linearization |
| arog 13s fw | taggtctcagcccgacgagggtatcatatgaattatcagaacgacg | aroG^D146N^ plasmid | aroG^D146N^ linearization |
| arog 13s rv | taggtctcatgacttacccgcgacgcgcttttactgc | aroG^D146N^ plasmid | aroG^D146N^ linearization |
| arob 13s fw | taggtctcagtcagacgagggtatcatatgcagacactaaaggtcgacctg | gDNA *P. putida* KT2440 | aroB linearization |
| arob 13s rv | taggtctcaccgttcaaagctgggccacgatcgc | gDNA *P. putida* KT2440 | aroB linearization |
| aroq 13s fw | taggtctcaacgggacgagggtatcatatggcaacgctactggtgctcc | gDNA *P. putida* KT2440 | aroQ linearization |
| aroq 13s rv | taggtctcagttctcatttgggctgtgcgttggcagc | gDNA *P. putida* KT2440 | aroQ linearization |
| aroe 13s fw | taggtctcagaacgacgagggtatcatatggaccagtacgtcgtttttgg | gDNA *P. putida* KT2440 | aroE linearization |
| aroe 13s rv | taggtctcaagtatcagccccgagcgagttg | gDNA *P. putida* KT2440 | aroE linearization |
| pabc 14wgib fw | ttaagattaactcacacacgaaagtatcatatgttcttaattaacggtcataagcag | gDNA *E. coli* K-12 | pabC linearization |
| pabc 14wgib rv | atgatactttcgtattcgggcgctcacaaag | gDNA *E. coli* K-12 | pabC linearization |
| aroq 14wgib fw | tttgtgagcgcccgaatacgaaagtatcatatggcaacgctactggtgctcc | gDNA *P. putida* KT2440 | aroQ linearization |
| aroq 14wgib rv | atgatactttcgttcatttgggctgtgcgttggcagcc | gDNA *P. putida* KT2440 | aroQ linearization |
| aroe 14wgib fw | acgcacagcccaaatgaacgaaagtatcatatggaccagtacgtcgtttttgg | gDNA *P. putida* KT2440 | aroE linearization |
| aroe 14wgib rv | atgatactttcgttcagccccgagcgagttg | gDNA *P. putida* KT2440 | aroE linearization |
| tyra 14wgib fw | aactcgctcggggctgaacgaaagtatcatgtggtaaatgcagcaaaaacc | gDNA *P. putida* KT2440 | tyrA linearization |
| tyra 14wgib rv | atgatactttcgttcacgacttgccctcttctgcc | gDNA *P. putida* KT2440 | tyrA linearization |
| aroc 14wgib fw | aagagggcaagtcgtgaacgaaagtatcatatgtccggcaatacctacggcaagc | gDNA *P. putida* KT2440 | aroC linearization |
| aroc 14wgi rv | ccttgcccttttttgccggatcagcgctggcccagcaccg | gDNA *P. putida* KT2440 | aroC linearization |
| paba 14s fw | taggtctcagcccgacgagggtatcatatgatcctgcttatagataactacg | gDNA *E. coli* K-12 | pabA linearization |
| paba 14s rv | gtggtctctttggtcagcgatgcaggaaattagcc | gDNA *E. coli* K-12 | pabA linearization |
| pabb 14s fw | taggtctcaccaagacgagggtatcatatgaagacgttatctcccgc | gDNA *E. coli* K-12 | pabB linearization |
| pabb 14s rv | gtggtctcttctgttacttctccagttgcttc | gDNA *E. coli* K-12 | pabB linearization |
| arog 14s fw | taggtctcacagagacgagggtatcatatgaattatcagaacgacg | aroG^D146N^ plasmid | aroG^D146N^ linearization |
| arog 14s rv | taggtctcatgacttacccgcgacgcgcttttactgc | aroG^D146N^ plasmid | aroG^D146N^ linearization |
| arob 14s fw | taggtctcagtcagacgagggtatcatatgcagacactaaaggtcgacctg | gDNA *P. putida* KT2440 | aroB linearization |
| arob 14s rv | taggtctcaccgttcaaagctgggccacgatcgc | gDNA *P. putida* KT2440 | aroB linearization |
| arok 14s fw | taggtctcaacgggacgagggtatcatgtgcgaaatttgatacttgtgg | gDNA *P. putida* KT2440 | aroK linearization |
| arok 14s rv | taggtctcaagtattaacggggcggcaactgctgc | gDNA *P. putida* KT2440 | aroK linearization |
| paba 15w fw | taggtctcagcccgacgaaagtatcatatgatcctgcttatagataactacg | gDNA *E. coli* K-12 | pabA linearization |
| paba 15w rv | gtggtctctttggtcagcgatgcaggaaattagcc | gDNA *E. coli* K-12 | pabA linearization |
| pabb 15w fw | taggtctcaccaagacgaaagtatcatatgaagacgttatctcccgc | gDNA *E. coli* K-12 | pabB linearization |
| pabb 15w rv | gtggtctctaccgttacttctccagttgcttc | gDNA *E. coli* K-12 | pabB linearization |
| aroq 15w fw | taggtctcacggtgacgaaagtatcatatggcaacgctactggtgctcc | gDNA *P. putida* KT2440 | aroQ linearization |
| aroq 15w rv | taggtctcagttctcatttgggctgtgcgttggcagc | gDNA *P. putida* KT2440 | aroQ linearization |
| aroe 15w fw | taggtctcagaacgacgaaagtatcatatggaccagtacgtcgtttttgg | gDNA *P. putida* KT2440 | aroE linearization |
| aroe 15w rv | taggtctcatgactcagccccgagcgagttg | gDNA *P. putida* KT2440 | aroE linearization |
| arok 15w fw | taggtctcagtcagacgaaagtatcatgtgcgaaatttgatacttgtgg | gDNA *P. putida* KT2440 | aroK linearization |
| arok 15w rv | taggtctcaagtattaacggggcggcaactgctgc | gDNA *P. putida* KT2440 | aroK linearization |
| aroc 15w fw | taggtctcatactgacgaaagtatcatatgtccggcaatacctacgg | gDNA *P. putida* KT2440 | aroC linearization |
| aroc 15w rv | aggtctcgactgcagcggccgctactagtattattatcagcgctggcccagcaccggggtattc | gDNA *P. putida* KT2440 | aroC linearization |
| pabc 15sgib fw | ttaagattaactcacacacgagggtatcatatgttcttaattaacggtcataagcag | gDNA *E. coli* K-12 | pabC linearization |
| pabc 15sgib rv | atgataccctcgtattcgggcgctcacaaag | gDNA *E. coli* K-12 | pabC linearization |
| arog 15sgib fw | tttgtgagcgcccgaatacgagggtatcatatgaattatcagaacgacg | aroG^D146N^ plasmid | aroG^D146N^ linearization |
| arog 15sgib rv | atgataccctcgtttacccgcgacgcgcttttactgc | aroG^D146N^ plasmid | aroG^D146N^ linearization |
| arob 15sgib fw | aagcgcgtcgcgggtaaacgagggtatcatatgcagacactaaaggtcgacctg | gDNA *P. putida* KT2440 | aroB linearization |
| arob 15sgib rv | atgataccctcgttcaaagctgggccacgatcgc | gDNA *P. putida* KT2440 | aroB linearization |
| tyra 15sgib fw | tcgtggcccagctttgaacgagggtatcatgtggtaaatgcagcaaaaacc | gDNA *P. putida* KT2440 | tyrA linearization |
| tyra 15sgib rv | ccttgcccttttttgccggatcacgacttgccctcttctg | gDNA *P. putida* KT2440 | tyrA linearization |
| paba 16sgib fw | ttaagattaactcacacacgagggtatcatatgatcctgcttatagataactacg | gDNA *E. coli* K-12 | pabA linearization |
| paba 16sgib rv | atgataccctcgttcagcgatgcaggaaattagcc | gDNA *E. coli* K-12 | pabA linearization |
| pabb 16sgib fw | atttcctgcatcgctgaacgagggtatcatatgaagacgttatctcccgc | gDNA *E. coli* K-12 | pabB linearization |
| pabb 16sgib rv | atgataccctcgtttacttctccagttgcttc | gDNA *E. coli* K-12 | pabB linearization |
| pabc 16sgib fw | agcaactggagaagtaaacgagggtatcatatgttcttaattaacggtcataagcag | gDNA *E. coli* K-12 | pabC linearization |
| pabc 16sgib rv | atgataccctcgtattcgggcgctcacaaag | gDNA *E. coli* K-12 | pabC linearization |
| arog 16sgib fw | tttgtgagcgcccgaatacgagggtatcatatgaattatcagaacgacg | aroG^D146N^ plasmid | aroG^D146N^ linearization |
| arog 16sgib rv | atgataccctcgtttacccgcgacgcgcttttactgc | aroG^D146N^ plasmid | aroG^D146N^ linearization |
| arob 16sgib fw | aagcgcgtcgcgggtaaacgagggtatcatatgcagacactaaaggtcgacctg | gDNA *P. putida* KT2440 | aroB linearization |
| arob 16sgib rv | atgataccctcgttcaaagctgggccacgatcgc | gDNA *P. putida* KT2440 | aroB linearization |
| aroq 16sgib fw | tcgtggcccagctttgaacgagggtatcatatggcaacgctactggtgctcc | gDNA *P. putida* KT2440 | aroQ linearization |
| aroq 16sgib rv | atgataccctcgttcatttgggctgtgcgttggcagcc | gDNA *P. putida* KT2440 | aroQ linearization |
| aroe 16sgib fw | acgcacagcccaaatgaacgagggtatcatatggaccagtacgtcgtttttgg | gDNA *P. putida* KT2440 | aroE linearization |
| aroe 16sgib rv | atgataccctcgttcagccccgagcgagttg | gDNA *P. putida* KT2440 | aroE linearization |
| arok 16sgib fw | aactcgctcggggctgaacgagggtatcatgtgcgaaatttgatacttgtgg | gDNA *P. putida* KT2440 | aroK linearization |
| arok 16sgib rv | atgataccctcgtttaacggggcggcaactgctgc | gDNA *P. putida* KT2440 | aroK linearization |
| tyra 16sgib fw | agttgccgccccgttaaacgagggtatcatgtggtaaatgcagcaaaaacc | gDNA *P. putida* KT2440 | tyrA linearization |
| tyra 16sgib rv | atgatactttcgttcacgacttgccctcttctgcc | gDNA *P. putida* KT2440 | tyrA linearization |
| aroc 16sgib fw | aagagggcaagtcgtgaacgaaagtatcatatgtccggcaatacctacggcaagc | gDNA *P. putida* KT2440 | aroC linearization |
| aroc 16sgib rv | ccttgcccttttttgccggatcagcgctggcccagcaccg | gDNA *P. putida* KT2440 | aroC linearization |
| Promoter | cagacctggaattgtgagc | Constructed strains *P. putida* KT2440 | Colony PCR |
| Pp | accttgcccttttttgccggtctcgactgcagcg | Constructed strains *P. putida* KT2440 | Colony PCR |

Figure S1. Genetic parts of each *P. putida* KT2440 strain S1-S16 constructed in this study

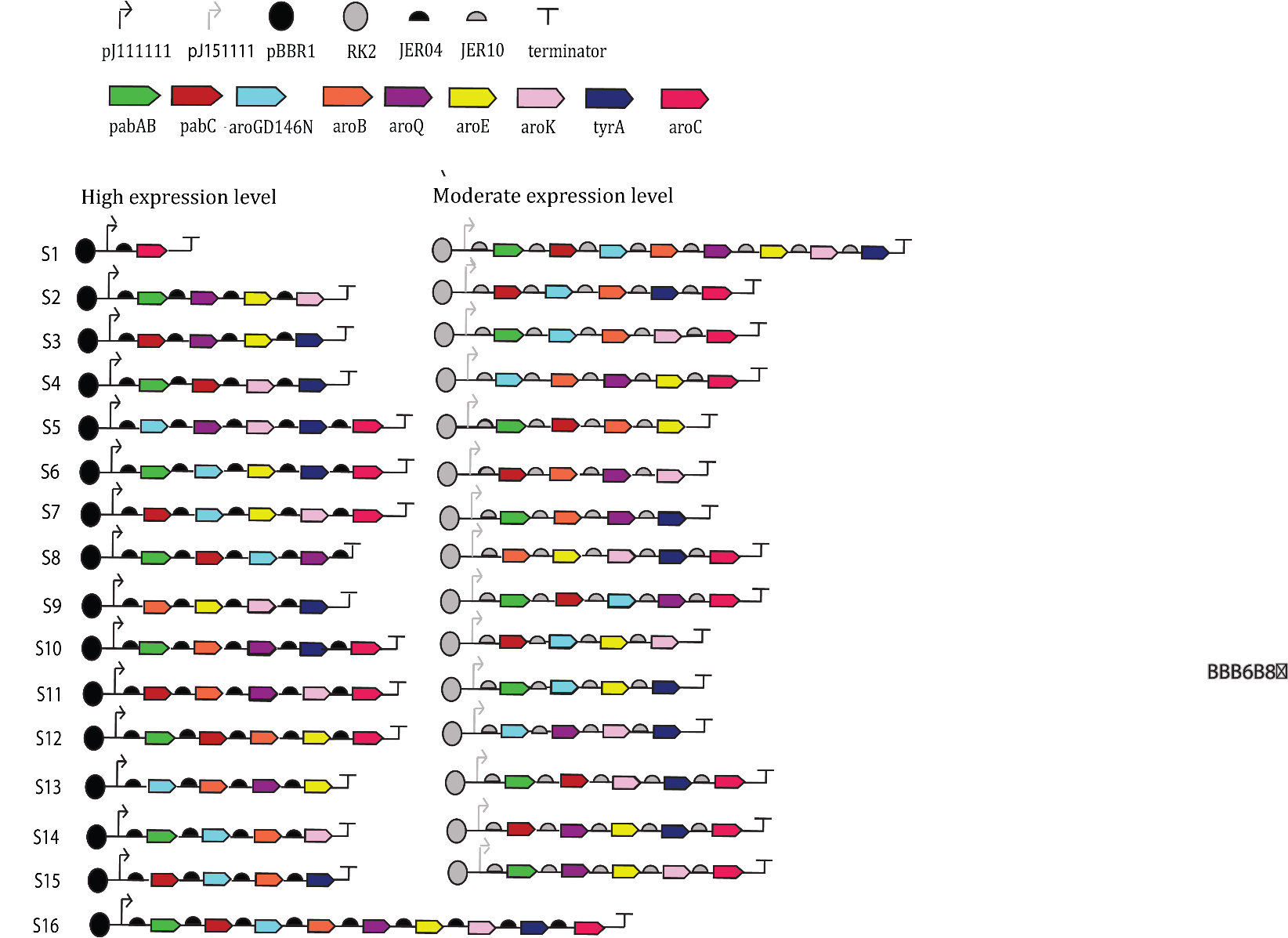
